# Impending anthropogenic threats and protected area prioritization for the largest Neotropical apex predator in its Amazonian stronghold

**DOI:** 10.1101/2021.08.31.458336

**Authors:** Juliano A. Bogoni, Valeria Boron, Carlos A. Peres, Maria Eduarda M. S. Coelho, Ronaldo G. Morato, Marcelo Oliveira-da-Costa

## Abstract

Jaguars (*Panthera onca*) exert critical top-down control over large vertebrates across the Neotropics and have been declining due to multiple threats. Based on geospatial layers, we extracted socio-environmental variables for 447 protected areas across the Brazilian Amazon to identify protected areas that merit short-term high-priority efforts to maximize jaguar persistence. Data were analyzed using descriptive statistics, structural equations and regression modeling. Our results reveal that areas containing the largest jaguar densities and estimated population sizes are precisely those confronting most anthropogenic threats. We reveal that jaguars in the world’s largest tropical forest biome are threatened by deforestation associated with anthropogenic fires, and subsequent establishment of pastures. We provide a shortlist of protected areas that should be prioritized for short-term jaguar conservation. The future predicament of jaguar populations can only be ensured if protected areas can be proofed against downgrading and downsizing geopolitical pressures and external anthropogenic threats.

## Introduction

Large carnivores like jaguars *Panthera onca* exert critical roles in maintaining ecosystem health and integrity [1–2]. However, their populations are rapidly declining, and are particularly vulnerable to extinction because they occur at low densities, have slow population growth rates, and require large areas containing a healthy prey base to survive [3–4]. Thus, their long-term population viability requires large scale conservation planning approaches that include protected areas networks and connectivity corridors [e.g. 5–6]. Virtually all large-bodied wild carnivorans species have experienced clear population declines worldwide [2, 7]. Given that large-bodied felids often need large areas of appropriate habitat due to their large home ranges [7–8], these apex predators are strongly susceptible to high mortality in areas densely settled by humans [7, 9], even being able to often use agroecosystems as either corridors or supplementary habitats in fragmented landscapes [10–12].

The jaguar is the world’s third largest felid, and the largest in the Americas [13]. Jaguars exert considerable vertebrate top-down control and have populated the imagination of people since pre-Colombian days. The jaguar is therefore considered an emblematic flagship and a keystone species [1, 14]. The species ranges from the southern USA [15] to Argentina and is considered “Near Threatened” according to the IUCN Red List [16], and “Vulnerable” in Brazil [17]. Jaguars feed on a broad range of large-bodied terrestrial, semi-aquatic, and aquatic prey [18], resulting in large spatial requirements and wide-ranging movements to meet their daily metabolic needs [19–20]. This pattern of space use tends to increase as habitat quality decreases, rendering these apex hyper-carnivores particularly vulnerable to habitat loss and fragmentation [10, 12]. Due to their large spatial requirements, jaguars are also valuable in conservation planning, ensuring that many other co-occurring species and high-quality habitat are also protected [21].

Despite the cultural and ecological importance of the iconic jaguar, the species only occupies *ca.* 50% of its historic range [16] and has been almost extirpated from heavily modified Brazilian biomes, such as the Atlantic Forest and the Caatinga [22]. The main threats to jaguar survival are habitat loss, human persecution, and decline of prey populations [16]. This large apex predator was presumed extinct in over half of over 1,000 mammal assemblages spanning the entire Neotropical realm [23]. The Amazon forest still holds high jaguar densities and ~67% of the total jaguar range (~9 million km^2^) with higher probabilities of survival (e.g. [4, 24–25]). The Brazilian Amazon forest comprises ~77% of the Pan-Amazon region of South America [26], making it a high priority stronghold for jaguar conservation.

Despite a large network of protected areas (hereafter, PAs), the Brazilian Amazon has been encroached by deforestation frontier expansion, powered by non-natural fires, mining, and roads [27, 28], making conservation priority-setting increasingly necessary [29]. Amazonian deforestation rates have recently accelerated, leading to a process of savannization of both fauna and flora throughout the so-called “deforestation arc” of the Brazilian Amazon [30, 31]. Annual deforestation in the Brazilian Amazon in 2018-2019, estimated at ~1,760,000 hectares, was further aggravated by unprecedented anthropogenic fire events [32, 33], with the peak deforestation year in a decade recorded in 2020 [34]. Under this complex socio-environmental context, the PA network across the Brazilian Amazon is crucial for jaguars and biodiversity conservation [35, 36]. Across the Brazilian Amazon, there are 307 federal and state-managed conservation units (UCs), 196 of which are sustainable use and 111 are strictly protected reserves, comprising 23.5% (~1.18 million km^2^) of Brazilian Amazonia. An additional 23% of Brazilian Amazonia (~1.16 million km^2^) is protected ‘on paper’ by 424 indigenous reserves (ILs), but their fate remains highly uncertain [37].

Conservation units and indigenous reserves are recognized as pivotal tools in retaining relatively intact biotas worldwide, and buffering tropical forest climate tipping-points by retaining aboveground carbon storage [38]. Considering the vast but severely underfunded network of PAs, the dilemma of prioritizing conservation investments in the short, medium, and long term is paramount for successful conservation outcomes [39]. A fine-tuned conservation plan for a focal species such as the jaguar can serve as a robust proxy for overall biodiversity persistence [21]. Similar study was developed across the Paleotropics to evaluate the performance of PAs in still harboring viable population of African lions (*Panthera leo*) and their basis of prey [7]. This range-wide assessment of PAs with African lions revealed multiple socio-environmental factors of threats, such as hunting, human and livestock encroachment, and human-wildlife conflicts [7]. Despite that, PAs within lion range have potential to host a cumulative population size that is up to four times larger than the current total free-ranging lion population, even Africa’s PA networks will continue to deteriorate, and the effectiveness depends on their legal status and management [7]. Across the Neotropics the jaguar population declines also coincides with similar human-induced pressures revealed to lions in Africa [see 12]. Given that PAs are central to safeguarding biodiversity, these protected lands are under multiple geopolitical pressures and their nominal buffer zones are highly degraded as the wider unprotected countryside [40].

Therefore, we aimed to (1) identify the main socio-environmental threats to jaguars across the Brazilian Amazon, (2) assess to what degree the main threats faced by jaguars are related to their population sizes within PAs, (3) assess whether the legal denomination of PAs affects jaguar population sizes and threats, and (4) identify PAs that merit high-priority short-term conservation action to safeguard these apex carnivores. Our hypotheses are that (1) habitat degradation factors, such as deforestation and wildfires, are the most important and imminent threats to jaguar survival across the Brazilian Amazon; (2) the legal denomination of PAs is an important determinant of PA threat status; and (3) PAs that should be prioritized for jaguar conservation efforts are precisely those confronting the most severe habitat degradation threats. Our study is important to better understand current threats to jaguars, helping to align conservation efforts by presenting an evidence-based agenda for jaguar conservation in the Amazon.

## Material and Methods

### Ethics Statement

This work was not submitted to an Institutional Animal Care and Use Committee (IACUC) or equivalent animal ethics committee, because the data were only based on spatial data-frames, polygons or raster files.

### Study meta-region: Brazilian Amazon

The Brazilian Amazon represents ~50% of Brazil’s territory and ≌76.8% (i.e. 5.15 million km^2^) of the ~7.59 million km^2^ Pan-Amazon, spanning nine South American countries. The Amazon region contains the vast majority of all Brazilian protected areas. However, the network of protected areas across the Brazilian Amazon is under increasing pressure linked to deforestation and other illegal activities [41–42]. This vast tropical biome is characterized mainly by tropical moist broadleaf forests [43] and has a human population within Brazil of ~23 million people, 72% of which living in major cities [44]. We scoped this study to include all officially sanctioned protected areas across the Brazilian Amazon, including 117 conservation units and 330 indigenous reserves, amounting to 1,755,637 km^2^ (S1 Fig), which represents 41.7% of Brazilian Amazonia. Among these 117 conservation units, 57 are strictly protected areas (SPAs) and 60 are sustainable use reserves (SURs). Among the 330 indigenous reserves, 37 have only been ‘declared’ as such (grouped as IR type IR1; lands that obtained the issuance of the Declaratory Ordinance by the Minister of Justice and are authorized to be physically demarcated, with the materialization of landmarks and georeferencing), whereas 293 have been physically demarcated and officially sanctioned (grouped as IR type IR2; Lands that have their boundaries materialized and georeferenced, whose administrative demarcation was ratified by a Presidential decree and/or lands that, after the ratification decree, were registered in the Notary Office in the name of the Union and in the Secretary of Heritage of the Union) (S1 File).

### Data acquisition

We used several high-resolution spatial layers (rasters or polygons) to extract variables for each protected area that represent (1) a proxy of direct jaguar mortality (e.g. roadkill and persecution due to livestock depletion; Michalski et al., 2006): (i) human population density (HPD), sourced from the Brazilian Institute of Geography and Statistics (spatial scale of 1:250,000) [45]; (ii) road density (include paved and unpaved), sourced from the Brazilian Institute of Geography and Statistics (spatial scale of 1:250,000) [46]; (iii) pasture area, sourced from MapBiomas (v.5, spatial resolution of 30 m) [47]; and layers that represent

(2) habitat degradation: (i) fire hotspots over a 5-yr period (2016 - 2020), sourced from the National Institute for Space Research-INPE (MODIS sensor by TERRA reference satellite; spatial resolution of 1 km) [32]; (ii) deforestation over 4 years (2016 - 2019) sourced from PRODES (spatial resolution of 30 m) [32]; and (iii) mining areas, sourced from MapBiomas (v.5, spatial resolution of 30 m) [47]. We also obtained the size of each protected area and its adjacent 5-km buffer zone.

Based on the SIRGAS-2000 UTM-ZONE 22°S projection, spatial data extraction was performed separately for both the internal PA area and the external 5-km buffer based on the administrative polygons of each protected area. We used a 5-km buffer threshold because this is approximately two-fold the minimum home range radius of Amazonian jaguars (i.e. 2.53 km for females, *ca.* 20 km^2^) [24]. Moreover, we sourced data on jaguar population density inside each protected area from Jędrzejewski et al. (2018) [25]. Due to the large pixel-size of jaguar population estimates (i.e., 10 km × 10 km; 100 km^2^) in Jędrzejewski et al. (2018) [25], we projected into each 5-km buffer the same average density obtained from each PA but arbitrarily penalized this density value by −40% due to a presumed “edge effect” [48] to avoid any overestimation of jaguar population size across the Brazilian Amazon. Data extraction was conducted using the ArcGIS 10.8 software [49] based on the average or sum of pixels/area both inside and outside each PA. Further, we obtained the type of legal denomination of each protected area (according to Sistema Nacional de Unidades de Conservação [SNUC; 50], and based on Ministério do Meio Ambiente [MMA; 51]) and the stage of legal implementation of each indigenous reserve (i.e. declared, approved, physically demarcated, and legally sanctioned) sourced from Fundação Nacional do Índio [52].

### Data analysis

#### Jaguar population responses to threats

We previously tested for differences among PA types (i.e., IR1, IR2, SPA, SUR) in how the main response variables (jaguar population size) responded to predictive variables (see below) using ANOVAs followed by Tukey post hoc comparisons by correcting for data asymmetry using log_10_ (x + 1) [53]. To disentangle the likely relationships between potential future drivers of jaguar population size both inside and outside PAs, we used path analysis (i.e., structural equation modeling [SEM]) [54]. SEM was implemented to simultaneously evaluate the relationship of adjusted jaguar population size inside and outside PAs as an additive function. For jaguar population size within PAs, we included the following predictive variables: (i) protected area type (neutral), (ii) pasture area inside and outside; (iii) extent of roads both inside and outside PAs; and (iv) HPD inside and outside PAs as proxies of direct mortality; (v) deforestation inside and outside; (vi) fire severity inside and outside; and (vii) mining activities inside or outside as proxies of habitat degradation. Thus, the SEM model as implemented based on linear relationships as follows: jaguar population size_(inside)_ ~ PA type + pasture_(inside)_ + roads_(inside)_ + HPD_(inside)_ + deforestation_(inside)_ + fire_(inside)_ + pasture_(outside)_ + roads_(outside)_ + HPD_(outside)_ + deforestation_(outside)_ + fire_(outside)_ + mining_(outside)_ & jaguar population size_(outside)_ ~ PA type + pasture_(outside)_ + roads_(outside)_ + HPD_(outside)_ + deforestation_(outside)_ + fire_(outside)_ + mining_(outside)_. For jaguar population size within adjacent buffer areas, the SEM model included the same variables for PA polygons. All predictive variables were tested for multicollinearity using variance inflation factors (VIF) [55]. Results are shown using the SEM standardized coefficient (ssc; effect size), and we obtained R^2^-values based on the proportion of variance explained by each predictor [56]. All data were previously log-transformed (log_10_ (x + 1)). Data analysis was performed using R 4.0.2 [57] based on the *sem* function in the *lavaan* package [58]. We also performed the SEM analysis using the data of jaguar population size outside PAs without the prior penalization by –40%. For those variables yielding the strongest SEM coefficients, we plotted both linear and smoothed functions to present their explicit relationships with jaguar population sizes.

### Threat index for protected area prioritization criteria

We constructed a “*threat index*” (*TI*) applied to each of the 447 protected areas using the above geospatial layers for both the PA (_*in*_) and the buffer polygons (_*out*_), which are weighted according to SEM standardized coefficient and R^2^. To do so, the TI incorporated the (1) ratio of mining threats (*min*), defined as the size of mining operations (km^2^) in relation to PA size (km^2^), (2) pasture area (*pas*) defined as the size of pasture areas both inside and outside PA in km^2^, (3) ratio of deforestation amount over a 4-yr time-series (*def*), based on the amount of cumulative deforestation (km^2^) in relation to PA size; (4) total length (km) of roads (*roa*) overlapping each PA; (5) density of fires (*fir*) defined as the fire frequency over the 5-yr time-series divided by the PA size; and (6) maximum human population density (*hpd*) for each polygon area. We thus assigned relative weights to these variables to compose the TI according to prior SEM results. We also relativized the threat index by the maximum value at any protected area, which therefore ranged from 0 to 1 by dividing any *TI*_*i*_ for the max *TI*_*i,j*_. The threat index was obtained given the following equation:

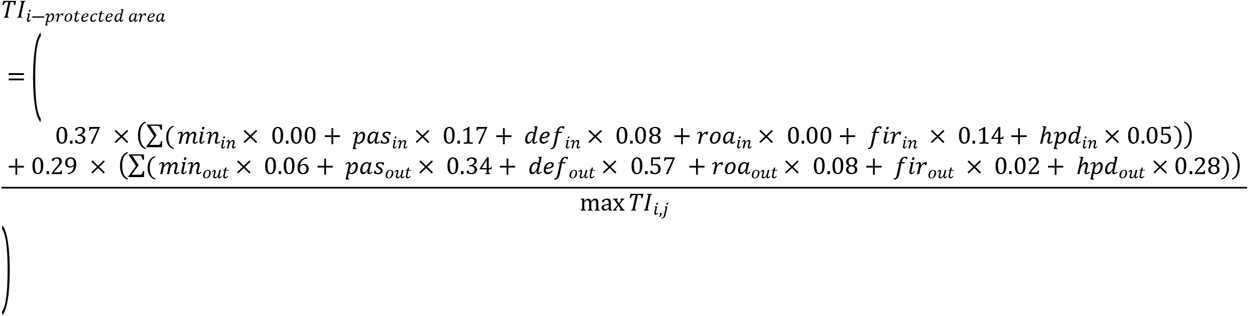

To identify PAs with the highest priority for short-term jaguar conservation action, we constructed bivariate plots between the (i) jaguar density inside any PA *vs.* the threat index; and (ii) jaguar population size inside any PA *vs.* the threat index. We used both jaguar density and total population size to avoid penalizing small PAs, since jaguar numbers are collinear with PA size. Based on the average of both variables in each bivariate plots, we defined one quadrant of short-time high priority based on: (1) high jaguar density and high threat index; and (2) large jaguar population and high threat index. We then identified the top-20 protected areas to be allocated conservation efforts across the Brazilian Amazon, selecting areas located in the extreme distribution across each quadrant by adding a tangential line within each quadrant that separates the top-10 areas. Once we identified the main spatial covariates related to jaguar population sizes, we then tested differences between high- and low-priority PAs using an ANOVA followed by Tukey post hoc tests while correcting for data asymmetry using log_10_ (x + 1) [53].

## Results

### Threats to jaguar across Amazonian protected areas

The potential jaguar density was on average 1.55 individuals per 100 km^2^ (± 0.75 *sd* ranging from 0.00 to 4.22 ind.) across the 447 protected areas in the Brazilian Amazon and their respective 5-km radial buffers, hereafter the wider PA area. This amounted to a combined area of 2,244,090 km^2^, which could support 34,784 (95% CIs: 26,922 - 42,644) jaguars. Between 2016 and 2019 deforestation across protected areas and their respective 5-km radial buffers amounted to 5,560 km^2^, representing 0.25% of the total area. There were also 101,804 agricultural and forest understorey fires over a 5-yr period, and roads across these protected areas and their buffers amounted to 3,947 km. We found significant differences in both jaguar population sizes and levels of threat among protected area types (Fig. 1; S2 File). While the largest jaguar populations were concentrated in strictly-protected (SPA) and sustainable-use reserves (SUR) compared to Indigenous Reserves, the threat index was higher in SURs than in SPAs and Indigenous Reserves (Fig. 1). None of the other variables showed significant differences among PA 0.57; p < 0.01) and inside (ssc = 0.66; p < 0.01) protected areas (Fig. 2). The extent of pasture areas and the HPD outside PAs were also related to number of jaguars inside and outside protected areas (pasture: ssc_jaguar inside_ = 0.44; p < 0.01; ssc_jaguar outside_ = 0.34; p < 0.01; HPD: ssc_jaguar inside_ = 0.28; p < 0.01; ssc_jaguar outside_ = 0.24; p < 0.01) (Fig. 4). Other variables related to jaguar population size were PA type (ssc_jaguar inside_ = 0.10; p = 0.02; ssc_jaguar outside_ = 0.10; p = 0.02), pasture area inside (ssc_jaguar inside_ = 0.19; p < 0.01), deforestation inside (ssc_jaguar inside_ = 0.08; p < 0.01), fires inside (ssc_jaguar inside_ = 0.14; p < 0.01), fires outside (ssc_jaguar inside_ = 0.17; p < 0.01), and total length of roads within buffer areas (ssc_jaguar outside_ = 0.08; p = 0.05; Fig. 2). The predictive power of the SEM model was R^2^ = 0.37 and R^2^ = 0.29 for jaguar populations inside and outside PAs, respectively (Fig. 2). The SEM model results, trends, coefficients, standard errors and covariance can be accessed in the S3 File. Further, the SEM analysis upon absolute values of jaguar population size outside the PA (i.e., without the 40% penalization) was exactly the same results aforementioned given that the only change was the order of magnitude of jaguar population size outside PAs, not the variation. Bivariate regression plots containing the main habitat disturbance predictors (i.e., deforestation, pastures and HPD), indicated that some relationships with jaguar population sizes were markedly nonlinear (Fig. 3) and the largest amounts of within-PA anthropogenic disturbance coincided with the ‘arc of deforestation’ region (Fig. 3G).

**Fig. 1.**
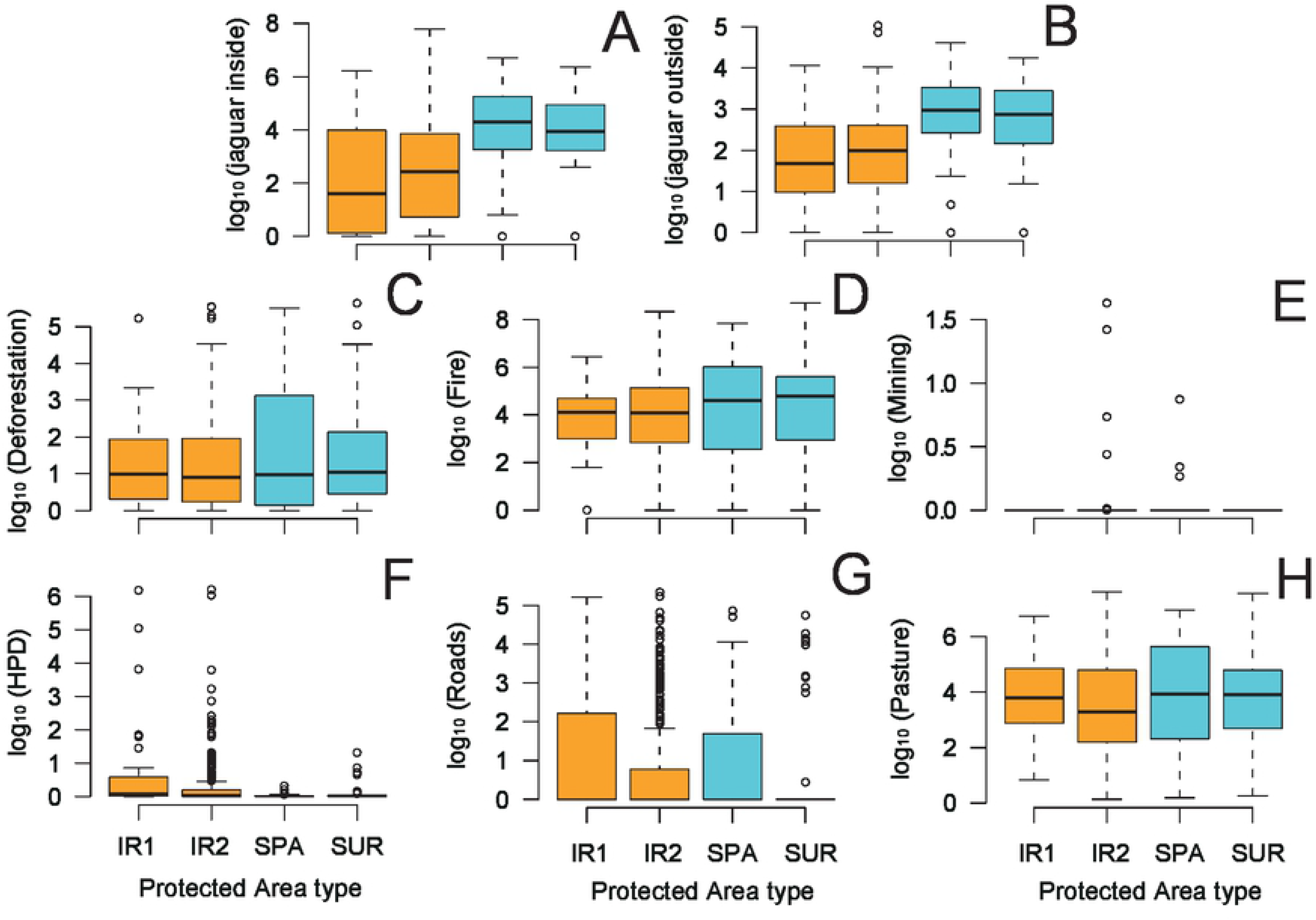
Jaguar population size inside protected areas (A), jaguar population size outside protected areas (5-km buffer; B), deforestation (km^2^; C), fire hotspots (N; D), mining areas inside (km^2^; E), human population density (ind/km^2^; F), roads (km, G), and exotic cattle pasture (km^2^, H) across 447 protected areas grouped by the legal status in the Brazilian Amazon. Acronyms are: SPA: Strictly Protected Conservation Units; SUR: Sustainable Use Conservation Units (SUR); IR1: Indigenous Reserves declared; and IR2: Indigenous Reserves demarcated, approved, or legally sanctioned. All data are log_10_-transformed and from C to H are the sum of values both inside and outside protected areas.

**Fig. 2.**
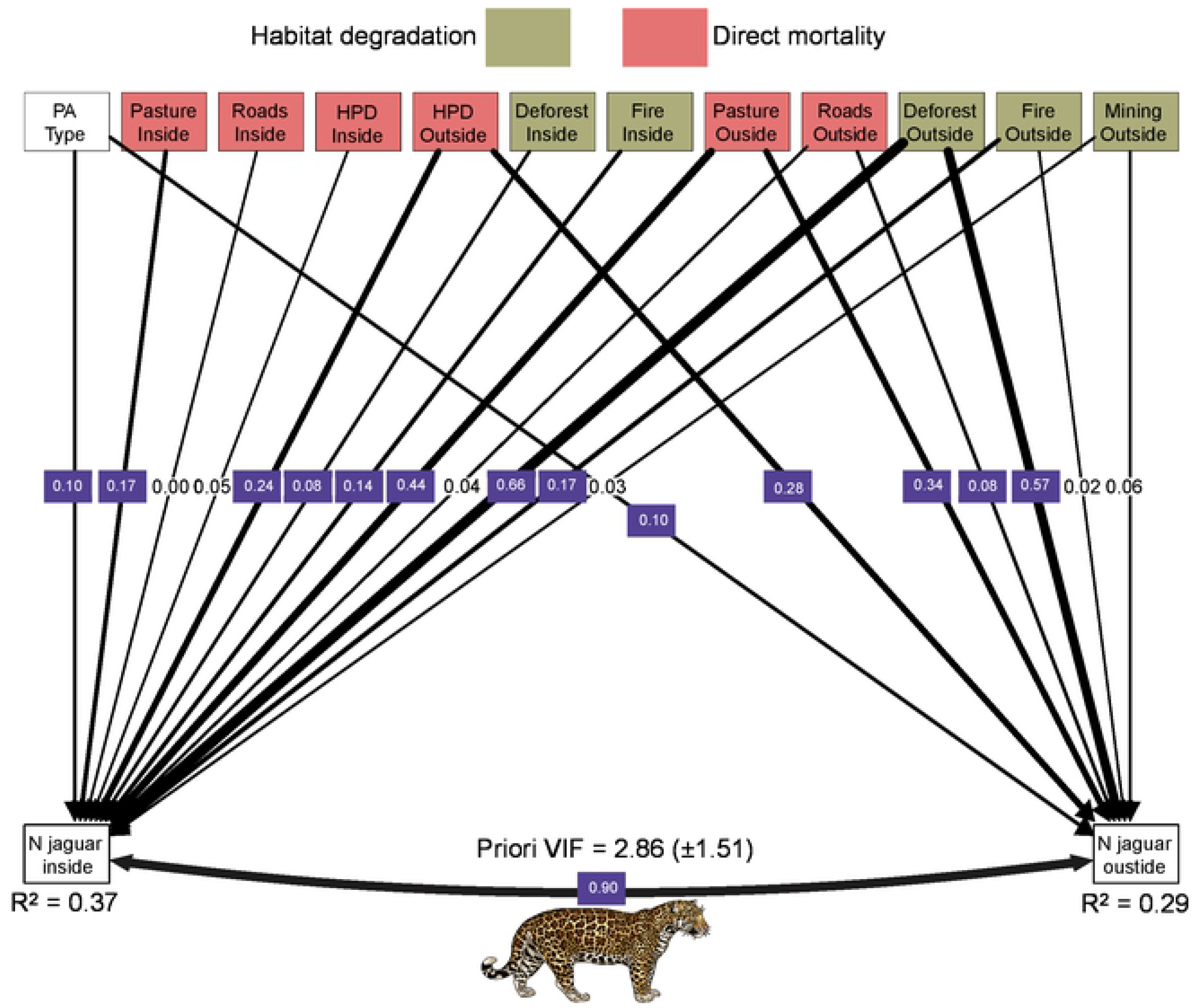
Structural equation models (SEM: path analysis) to disentangle the linear relationships between different environmental and demographic variables and either the Jaguar population size inside and outside (5-km radial buffer) across 447 protected areas located at the Brazilian Amazon. All data are presented in log_10_-transformation. Significant relationships (p < 0.05) are highlighted in purple.

**Fig. 3.**
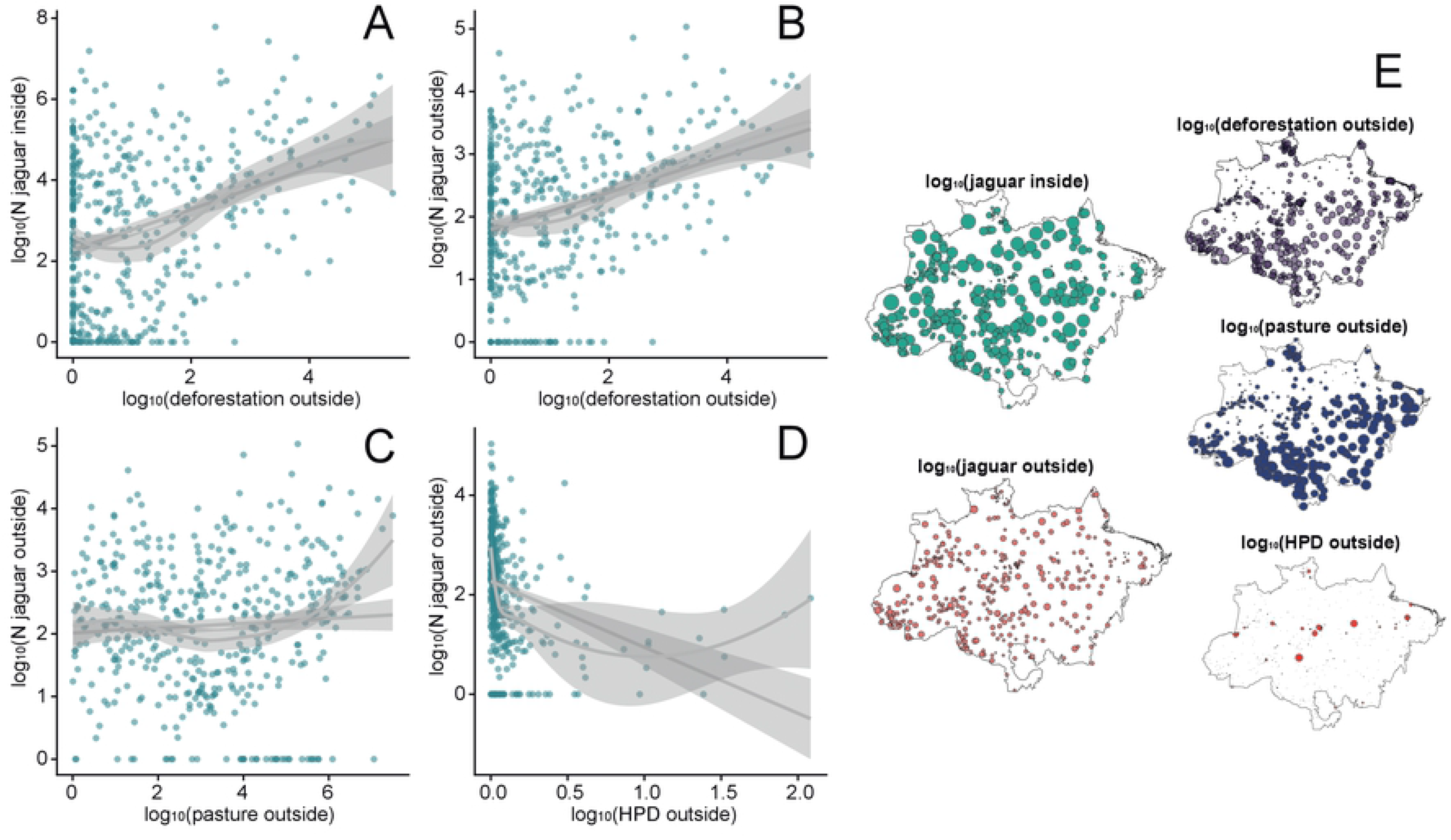
Bivariate plots and linear/non-linear relationships (and 95% CIs) between (A) jaguar population size inside and deforestation in 5-km radial buffer outside 447 protected areas located in the Brazilian Amazon; (B) Jaguar population size outside and deforestation outside; (C) Jaguar population size outside and pasture amount outside; (D) Jaguar population size outside and human population density; and the spatial distribution of these variables (E). All data are log_10_-transformed.

**Fig. 4.**
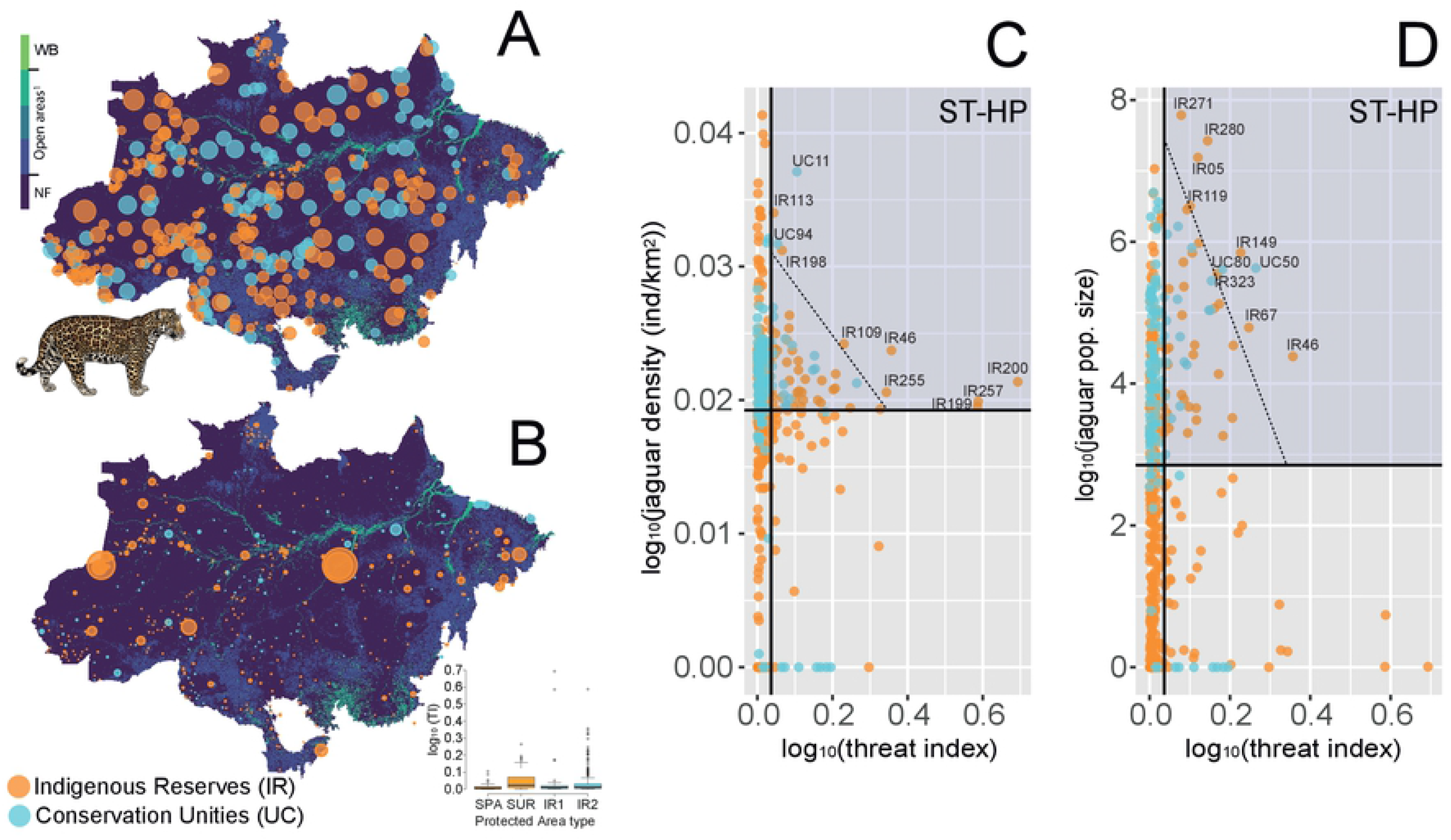
Distribution of jaguar population density (log_10_ x) inside protected areas (A), threat index and the values per protected area type (B) and bivariate plots between threat index and jaguar density inside protected areas (C), and between threat index and jaguar population size inside protected areas (D) across 447 protected areas in the Brazilian Amazon. Acronyms are ST-HP: short-term high priority quadrant (delimited by highlighted gray-frame) and the respective top-10 areas that should be prioritized in each approach based on the extreme of distributions threshold by a tangential line. We also identified 43 (9.7%) additional Amazonian PAs that should be prioritized for jaguar conservation in the short to medium term according to our prioritization quadrants (highlighted gray-frames). The background map in A and B is the land cover and land use dated from 2019 sourced at MapBiomas project (https://mapbiomas.org/) [47]. Map acronyms are WB: water bodies; NF: natural forest. ^1^Open areas include non-forest natural formation, farming, and non-vegetation area. For all land cover and land use classes see MapBiomas project [47].

### Protected area prioritization

Our proposed threat index was on average TI = 0.04 (± 0.093), but higher threat values were concentrated in PAs located within or near agricultural and logging frontiers in the southern Amazon (Fig. 4B). Based on the threat index, jaguar density, and jaguar population size of each PA, we were able to identify 19 non-redundant protected areas (4.25%) that deserve urgent high-priority conservation attention (Fig. 4C and 4D; Fig. 5; Table 1). These areas amount to a total of 398,228 km^2^ (22.7% of the aggregate wider PA area), and can support an estimated 7,843 jaguars under an average threat index of TI = 0.298 (± 0.278). Further, these 19 PAs experienced cumulative deforestation over 4 years of 259.1 km^2^ (5.0% of the total amount of deforestation within PAs amounting to 5,560 km^2^), 8,090 cumulative fire hotspots over 5 years (7.9% of all 101,804 fires), contain 105.1 km of roads (2.7% of the total road network within PAs and buffer zones of 3,947 km), 4,168 km^2^ of pastures (8.7% of 47,936 km^2^ of pastures within PAs), and an average HPD of 33.46 (± 113.6) people living within all wider PA areas (HPD = 2.14 ± 27.46 people/km^2^). These areas — comprising 15 indigenous reserves and four conservation units — are located near the Amazonian ‘arc of deforestation’ and northern Amazon (Fig. 5A; Table 1). These 19 PAs presented significantly higher deforestation rates (F = 4.70; p = 0.03), marginally more severe or more frequent fires (F = 2.88; p = 0.09), higher HPDs (F = 12.43; p < 0.001), but did not differ in pasture areas (F = 1.84; p = 0.18) compared to other protected areas (Fig. 5). We also identified 43 (9.7%) additional Amazonian PAs that should be prioritized for jaguar conservation in the short to medium term according to our prioritization quadrants (Fig. 4C and 4D).

**Fig. 5.**
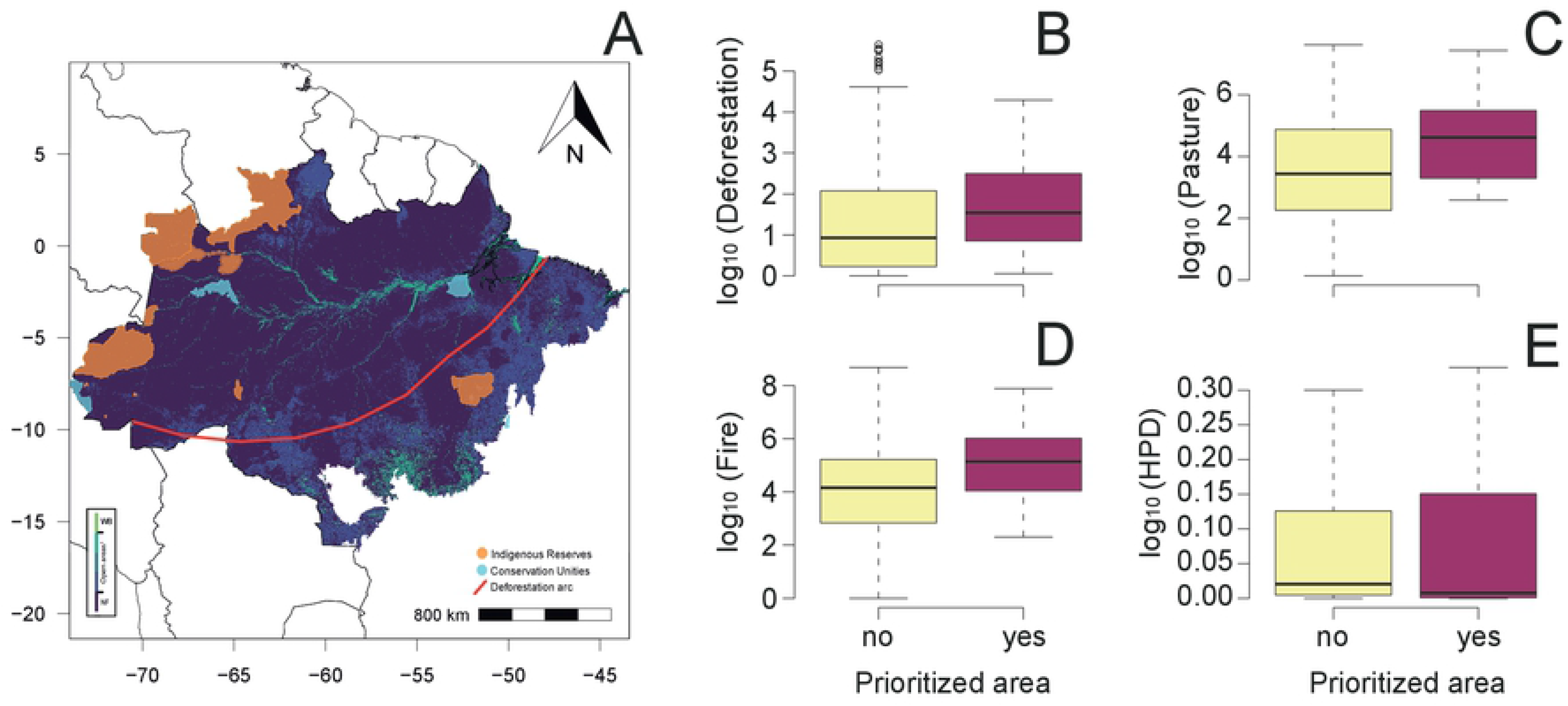
(A) Location of 19 protected areas in the Brazilian Amazon prioritized for jaguar conservation considering threat index, jaguar density, and jaguar population size. The background map is the land cover and land use dated from 2019 as sourced from the MapBiomas project (https://mapbiomas.org/) [47]. Map acronyms are WB: water bodies; NF: natural forest. ^1^Open areas include non-forest natural vegetation, farming, and non-vegetation areas. For all land cover and land use classes see MapBiomas project [47]. (B) Deforestation (km^2^); (C) Pasture (km^2^); (D) Fire focus (N of focus) comparison between areas that should be prioritized for Jaguar conservation across the Brazilian Amazon; and (E) Human population density (HPD) comparison between areas that should be prioritized for jaguar conservation across the Brazilian Amazon. In B, C, D and E the data are presented in log_10_-transformation.

**Table 1.** Protected area codes, names, size (km^2^), and legal status of reserves that should be highly prioritized for jaguar conservation across the Brazilian Amazon. Acronyms are SPA: Strictly Protected Conservation Units; SUR: Sustainable Use Conservation Units (SUR); IR1: Indigenous Reserves declared; IR2: Indigenous Reserves delimited, approved, or regularized; and TI: threat index. See the prioritization approach at Fig. 4, among top-20 protected areas we have 19 non-redundant protected area that should be prioritized at short-term (see Fig. 5).

| Code | Protected area name | Area (km<sup>2</sup>) | Status | Criteria |
| --- | --- | --- | --- | --- |
| IR199 | *Praia do Índio* | 0.3 | IR1 | High TI & high jaguar density |
| IR200 | *Praia do Mangue* | 0.3 | IR1 | High TI & high jaguar density |
| IR323 | *Jurubaxi-téa* | 12844.3 | IR1 | High TI & high jaguar pop. size |
| IR05 | *Alto Rio Negro* | 88019.5 | IR2 | High TI & high jaguar pop. size |
| IR46 | *Caititu* | 3289.2 | IR2 | Both |
| IR67 | *Évare I* | 6103.5 | IR2 | High TI & high jaguar pop. size |
| IR109 | *Katukina/Kaxinawá* | 260.6 | IR2 | High TI & high jaguar density |
| IR113 | *Kaxinawá do Baixo Rio Jordão* | 99.6 | IR2 | High TI & high jaguar density |
| IR119 | *Kayapó* | 32807.4 | IR2 | High TI & high jaguar pop. size |
| IR149 | *Médio Rio Negro I* | 19348.3 | IR2 | High TI & high jaguar pop. size |
| IR198 | *Poyanawa* | 284.5 | IR2 | High TI & high jaguar density |
| IR255 | *Tikuna de Santo Antonio* | 11.9 | IR2 | High TI & high jaguar density |
| IR257 | *Tukuna Umariáçu* | 54.2 | IR2 | High TI & high jaguar density |
| IR271 | *Vale do Javari* | 97013.5 | IR2 | High TI & high jaguar pop. size |
| IR280 | *Yanomami* | 100303 | IR2 | High TI & high jaguar pop. size |
| UC11 | *Parque Nacional da Serra do Divisor* | 9760.8 | SPA | High TI & high jaguar density |
| UC94 | *Parque Estadual do Cantão* | 1003.6 | SPA | High TI & high jaguar density |
| UC50 | *Reserva Extrativista Verde para Sempre* | 12894.3 | SUR | High TI & high jaguar pop. size |
| UC80 | *Reserva de Desenvolvimento Sustentável Mamirauá* | 14129.1 | SUR | High TI & high jaguar pop. size |

## Discussion

The persistence of healthy jaguar populations across their Amazonian stronghold depends on the implementation of public policies that safeguard the network of protected areas and their respective buffer zones. Since jaguars are classed as a flagship, umbrella [21] and a keystone species [59], the entire Amazonian biota can benefit from conservation efforts focused on this large felid. Our results reveal that the areas with the highest jaguar densities and largest estimated population sizes are precisely those most pressured by anthropogenic impacts in terms of habitat degradation. Deforestation, agricultural expansion including cattle pastures, and fires are prevalent in protected areas hosting the largest estimated jaguar populations, especially within their buffer areas, which fare far worse than their adjacent PAs [40]. As a large-bodied apex-predator, the large home range requirements of jaguars [19] frequently expose them to the edges and buffer zones of protected areas, coinciding with sites experiencing the most severe levels of habitat degradation. This contagious “edge effect” could determine jaguar numbers inside protected areas, by increasing mortality rates through, for example, shootings and roadkills, and the perverse effects of habitat fragmentation induced by deforestation [48]. On the other hand, our results also indicate that the network of PAs across the Brazilian Amazon which is the most important for jaguar conservation across the whole jaguar range is still fulfilling its role.

Across the Neotropics, drivers of local biotic depletion have accelerated since the 1970s. Dominant anthropogenic disturbances that lead to species declines and local extinctions include access to hitherto isolated forested areas via new roads [60], wildfires fueled by climate change [61], deforestation due to agribusiness frontiers expansion [62], relaxation of environmental law enforcement [63], increasing hunting pressure [64], and synergistic combinations between these and other socioeconomic stressors [23]. The Brazilian Amazon experienced an overwhelming and multifaceted spike in environmental degradation over the last decade, exacerbated by a renewed acceleration in deforestation and human-induced fires [34], exerting further pressure on Amazonian forest wildlife and native ethnic groups [37, 65].

Our results revealed that the relationship between fires and jaguar population sizes was prominently non-linear, indicating large jaguar populations at sites with both low and high fire severity, but particularly in frequently burned and fire-prone sites (see Fig. 3E). Typically, fire events represent the “*coup de grace*” following forest degradation, particularly where soil hydrological deficits are exacerbated. As powerful drivers of habitat degradation, deforestation and fires are historically synergistic, exerting a double-negative effect on tropical forest biotas [66]. Amazonian surface wildfires trigger a cascade of detrimental effects on biodiversity, particularly in areas that experienced little or no fire-stress over evolutionary timescales, leading to wholesale changes in species turnover [67]. It is therefore concerning that fire events are concentrated in areas containing large populations of jaguars and many other vertebrates. Our analysis also showed the potential risk of livestock pastures, which had also proliferated in areas packing large jaguar populations. Cattle ranches in the Amazon represent a double-whammy for jaguars. First, exotic pastures result directly from habitat loss for forest wildlife and, secondly, pasture-dominated landscapes become demographic sinks for jaguars, where as many as 110–150 large felids can be killed annually within a single Amazonian county through poisoned carcasses and direct persecution by professional jaguar/puma hunters and ranch staff [68].

Previous studies in human-modified landscapes showed that habitat loss and fragmentation have a strong detrimental impact on jaguar populations, which are now locally extinct in several Neotropical ecoregions [22–23]. For instance, the few remaining jaguar subpopulations in the Atlantic Forest are small, dispersed, and highly isolated into a few sufficiently large forest remnants [22, 69]. Our evidence indicates that consolidated Amazonian deforestation frontiers could exhibit similar patterns of jaguar demography within a few decades. For instance, jaguar populations potentially declined due the deforestation and wild fires by 1.8% in the last five years [70]. As jaguars have large spatial requirements, home range sizes and population density depend heavily on habitat quality and the prey base [12, 19]. In addition, jaguars strongly avoid non-forest areas in highly-forested landscapes [71]. Deforestation can also lead to further environmental degradation, including mining, roads and overhunting [72], increasing the threat to jaguars across their largest forest stronghold.

An additional factor of direct jaguar mortality is the expanding road network, particularly within areas surrounding PAs. Although the road network across the Amazon is still incipient, new major road projects can rapidly change this scenario and result in roadkills, and further habitat degradation, deforestation, and access by hunters to previously fremote areas [73–74]. The ubiquitous presence of road networks causes negative effects on mammal populations up to 5-km [75], strongly affecting apex predators across the tropics [e.g. 76]. In other Neotropical biomes, such as Atlantic Forest and the Cerrado, roadkills are an important contributor to jaguar mortality, further removing individuals from already depleted populations [22]. Thus, new government plans to expand the road network across the Amazon are an additional threat for jaguars and other wide-ranging carnivores within and around protected areas, which merits rethinking the strategic deployment of new infrastructure both for people and the environment [74].

These threat patterns are common to other apex-predators [e.g. 7, 77]. For instance, tiger populations (*Panthera tigris*) are highly threatened by recurrent forest loss across Indochina [77]. Although tigers are habitat generalists, most populations avoiding heavily disturbed areas and currently only survive in forest ecosystems, and core breeding populations are restricted to protected areas across much of their original range [77]. Accordingly, like in other wide-ranging large carnivores, a comprehensive network of effective protected areas throughout the Amazon is critical for the persistent of jaguar populations [e.g., 6, 76].

The largest jaguar populations were in strictly protected conservation units (SPAs), followed by multiple-use reserves (SUR) and legally demarcated and sanctioned Indigenous Reserves (IR2), whereas SURs were exposed to significantly higher levels of threat. However, individual threat variables comprising our threat metric did not vary between reserve denomination types (see Fig. 1 and S2 File). The 4-yr deforestation time-series reached 0.25% (5,560 km^2^) of the combined acreage across all protected areas and their respective buffers, an area 3.7-fold the size of São Paulo, the largest Latin American city. Moreover, cumulative fires over 5-yrs in the same combined area exceeded 100,000 burn hotspots. Other threats such as mining, road expansion, cattle pastures, and growing non-indigenous populations are also increasing across the Brazilian Amazon due to greater agricultural investments and relaxation in environmental legislation. This also shows that, through sheer lack of conservation investments, Amazonian nature reserves are beginning to fail their biodiversity conservation mission statements, much like early models of reserve defensibility predicted [78], and this predicament is often worse for indigenous territories. This study is also a warning to policy-makers as the situation will likely become worse in the near future, unless Amazonian PAs can be effectively protected.

Brazil’s state-managed protected areas are grounded in strict legislation under the National System of Conservation Units [see 50], but over the past few years, elevated geopolitical pressures have weakened the conservation assurances of these areas. Indigenous Reserves (also known as Indian Lands) are officially recognized to secure territorial rights for indigenous peoples and their traditional cultures, but withhold no legal property ownership (whether private or communal) over their own lands given that they are still demarcated as public lands [37]. Across the 447 protected areas examined here, indigenous reserves are presumed to contain ~21,700 jaguars, representing 62.4% of the total estimated number of jaguars across the Brazilian Amazon’s ~224 million hectares of protected areas and buffer zones. The importance of indigenous reserves is intrinsically linked to their larger sizes and larger wildlife populations compared to most state reserves, while ensuring legitimate land claims for native Amazonians as their original landholders [79].

Conservation priority-setting exercises are highly context-dependent in terms of socioeconomic dimensions [29] and prioritization dilemmas apply to poorly implemented protected areas that deserve urgent attention. Conservation efforts are typically limited by financial resources and economic models that assign priorities to these efforts have gained increasing importance [80]. We showed that our proposed threat metric could be an important priority-setting tool, and we were able to identify 16 protected areas that deserve immediate conservation efforts in the interest of long-term jaguar persistence. Interestingly, this approach identified that the geographic distribution of this set of protected areas is highly congruent with the Amazonian ‘arc of deforestation’. This region hosts the world’s largest mechanized agriculture frontier and includes the transitional ecotone between Amazonian forests and the Cerrado wooded scrublands, encompassing the Brazilian states of Maranhão, Tocantins, Pará, Mato Grosso, Rondônia, Acre, and more recently Amazonas. Chronic deforestation across the vast deforestation arc facilitates both human-wildlife conflicts between landowners and large felids, within which the latter always fares much worse [68]. This also calls for a more detailed analysis on the source-sink demographics of large cats in increasingly deforested hyper-fragmented landscapes that typically set reproductive viability thresholds for apex carnivore populations, such as the Harpy Eagle [81].

We recognize that our threat index can include biases on decision-making, which typically is a process of selecting between a wide range of management strategies based on uncertainty and incomplete information [82]. Nevertheless, the 19 non-redundant protected areas selected using this approach share a common potpourri of conservation troubles, including significant human population pressures and elevated levels of deforestation and wildfires. Further, we find significant differences in the main human-induced drivers of habitat degradation between protected areas that should be prioritized compared to PAs elsewhere. These 19 protected areas contained a 15-fold larger average human population size — but also a high *sd* — compared to other PAs, and typically are geographically positioned in the most pressured regions of the Amazon. Human population density is widely used to anchor geographic gradients of human disturbance and is a powerful proxy of human threats on natural ecosystems, including accessibility, infrastructure, land-use change, and direct mortality [5, 83].

Among the 19 protected areas assigned as urgent high priorities for immediate jaguar conservation effort across the Brazilian Amazon, we can highlight the strictly-protected 9,760.8-km^2^ *Parque Nacional da Serra do Divisor* near the Peruvian border. There has been heavy recent political pressure to build a road that would sever this park and link soybean production areas in Brazil with the Pacific. Across the Northwest Amazon, for instance, the *Vale do Javari* Indigenous Reserve — encompassing 97,014 km^2^ of nearly intact forest, and harboring the *Kulina Páno, Matis, Matses* ethnic groups — contains the largest estimated jaguar population (2,407 individuals) of the 477 protected areas examined here. However, this indigenous reserve faces a threat index of 0.19 fueled by deforestation, timber extraction, fires, and growing human populations, and more than double the average TI value across all PAs. The vast majority of Brazilian PAs, particularly in the Amazon, confront an average financial insufficiency of 89.7% [84]. In fact, this severe underfunding, understaffing and lack of operational infrastructure persists since the early 1990s [78] despite hundreds of millions of conservation dollars flowing into Brazil to boost the effectiveness of Amazonian PAs. Currently, the Brazilian government invests less than one dollar per km^2^ across all protected areas under state and federal jurisdiction. This is even more pronounced in the buffer zones surrounding conservation units [40], exacerbating jaguar population declines and weakening connectivity. Moreover, we highlight that the other 431 protected areas considered here do not necessarily deserve fewer conservation investments. For instance, 96 (21.5%) of all Amazonian PAs were classified as short to medium term priority as they face above-average levels of threat and hold above-average jaguar densities. We further acknowledge that the jaguar density estimate predictions that we used [25] do not include prey abundance data and can be biased. Thus, we recommend that future estimates of large felid population densities should include a measure of prey productivity. Overall, we reinforce that the threats faced by jaguars across the Brazilian Amazon consistently match the broad geographic patterns of habitat degradation.

Finally, the initial hypotheses we posed here were corroborated. Leading drivers of habitat degradation (i.e., deforestation and fires) are impending threats to large numbers of jaguars across the southern and eastern Amazon, and burgeoning human populations along an expanding agricultural frontier will likely inflict further mortality on both residents and transients. Further, other leading drives such as pasture area and HPD are confronting with PAs that harbor higher density or population size of jaguars. The most legally secure protected areas with the most restrictive use hosted the largest fraction of the overall estimated jaguar population, and reserves that should be prioritized for jaguar conservation are located at the epicenter of recent deforestation frontiers. We conclude that the main challenges faced by large carnivore conservation in the Amazon are deforestation associated with increasingly frequent and more severe anthropogenic fires. Using a snapshot of threat factors, we also provided a short-list of protected areas that deserves immediate conservation attention for jaguars and all co-occurring forest biodiversity. Currently, even in the most intact Neotropical regions, such as the Amazon and the Pantanal wetlands, the future of jaguars can be reasonably secure only in protected areas where land-use restrictions can be strictly enforced and that can resist the relentless political whims to downsize, downgrade and degazette them. *De facto* law enforcement, as opposed to protection “on paper” only, of healthy Amazonian ecosystems and their apex predators, will therefore require much greater political commitment than we have recently witnessed.

## Acknowledgements

We thank WWF network for the financial support. We thank to Luísa G. L. das Chagas for the support on spatial data extraction. JAB was supported by the São Paulo Research Foundation (FAPESP) postdoctoral fellowship grants 2018-05970-1 and 2019-11901-5.

## Supporting Information

**S1 Fig. Spatial distribution of 447 protected areas used to evaluate the threat to Jaguar across the Brazilian Amazon.**

**S1 File. Sociopolitical details of 447 protected areas used to evaluate the threat to Jaguar across the Brazilian Amazon.** Acronyms are: SPA: Conservation Unities of integral protection; SUR: Conservation Unities of sustainable use (SUR); IR1: Indigenous Reserves declared; and IR2: Indigenous Reserves delimited, approved, or regularized.

**S2 File. ANOVA and post hoc Tukey results comparing socioenvironmental variables between type of protected areas across the Brazilian Amazon.** SPA: Conservation Unities of integral protection; SUR: Conservation Unities of sustainable use (SUR); IR1: Indigenous Reserves declared; and IR2: Indigenous Reserves delimited, approved, or regularized.

**S3 File. SEM model results, trends, coefficients, standard errors and covariance in predicting jaguar population size inside and outside 447 protected areas across the Brazilian Amazon.**

## References

1. Estes J, Terborgh J, Brashares J, Power M, Berger J, Bond W, Carpenter S, Essington T, Holt R, Pikitch EK, Marquis R, Oksanen L, Oksanen T, Paine R, Ripple W, Sandin S, Scheffer M, Schoener T, Shurin J, Soule M, Virtanen R, Wardle D. Trophic downgrading of planet Earth. Science. 2011; 333: 301–306

2. Ripple WJ, Estes JA, Beschta RL, Wilmers CC, Ritchie EG, Hebblewhite M, Berger J, Elmhagem B, Letnic M, Nelson MP, Schmitz OJ, Smith DW, Wallach AD, Wirsing AJ. Status and Ecological Effects of the World’s Largest Carnivores Science. 2014; 343: 151–162.

3. Cardillo M, Mace GM, Jones KE, Bielby J, Bininda-Emonds ORP, Sechrest W, Orme CDL, Purvis A. Multiple causes of high extinction risk in large mammal species Science. 2005; 309: 1239–1241.

4. De La Torre JA, González-Maya JF, Zarza H, Ceballos G, Medellín RA. The jaguar's spots are darker than they appear: assessing the global conservation status of the jaguar (*Panthera onca*) Oryx 2018; 52(2): 300–315.

5. Sanderson EW, Redford KH, Chetkiewicz CLB, Medellin RA, Rabinowitz AR, Robinson JG, Taber AB. Planning to save a species: the jaguar as a model. Conservation Biology, 2002; 16(1): 58–72.

6. Rabinowitz, A, Zeller, KA. A range-wide model of landscape connectivity and conservation for the jaguar, *Panthera onca*. Biological Conservation 2010; 143: 939–945.

7. Lindsey PA, Pettracca LS, Funston PJ, Bauer H, et al. The performance of African protected areas for lions and their prey. Biological Conservation 2017; 209: 137–149.

8. Carbone C, Cowlishaw G, Isaac NJB, Rowcliffe JM How far do animals go? Determinants of day range in mammals. The American Naturalist 2005; 165: 290–297.

9. Woodroffe R. Predators and people: using human densities to interpret declines of large carnivores. Animal Conservation 2000; 3: 165–173.

10. Crooks, KR. Relative sensitivities of mammalian carnivores to habitat fragmentation. Conservation Biology 2002; 16: 488–502.

11. Ferreira AS, Peres CA, Bogoni JA, Cassano CG. Use of agroecosystem matrix habitats by mammalian carnivores (Carnivora): a global-scale analysis. Mammal Review, 2018; doi: 101111/mam12137.

12. Thompson JJ, Morato RG, Niebuhr BB, Bejarano V, et al. Range-wide factors shaping space use and movements by the Neotropic’s flagship predator: the jaguar Current Biology; 2021. https://doiorg/101016/jcub202106029.

13. Sunquist M, Sunquist F. Wild Cats of the World University of Chicago Press, Chicago, USA. 2002.

14. Leader-Williams N, Dublin HT. Charismatic megafauna as ‘flagship species’ In: Entwistle, A, Dunstone, N (eds) Priorities for the conservation of mammalian diversity: Has the panda had its day? Cambridge University Press, Cambridge pp 53−81. 2000.

15. Sanderson EW, Fisher K, Peters R, Beckmann JP, Bird B, Bradley CM, Bravo JC, Grigione MM, Hatten JR, González CAL, Menke K. A systematic review of potential habitat suitability for the jaguar Panthera onca in central Arizona and New Mexico, USA. Oryx, 2021; 1–12.

16. Quigley H, Foster R, Petracca L, Payan E, Salom R, Harmsen B. Panthera onca. The IUCN Red List of Threatened Species 2017: eT15953A50658693. 2017.102305/IUCNUK2017-3RLTST15953A50658693en Downloaded on 1 December 2020

17. Morato RG, Beisiegel BM, Ramalho EE, Boulhosa RLP. Avaliação do risco de extinção da Onça-pintada *Panthera onca* (Linnaeus, 1758) no Brasil. Biodiversidade Brasileira 2013; 3(1): 122–132.

18. Hunter L. Carnivores of the World Princeton Univ Press, Princeton, NJ. 2011.

19. Morato RG, Stabach JA, Fleming CH, Calabrese JM, De Paula RC, et al. Space Use and Movement of a Neotropical Top Predator: The Endangered Jaguar PLoS ONE 2016; 11(12): e0168176.

20. Chapman B, Hulthen K, Wellenreuther M, Hansson L-A, Nilsson JÅ, Bronmark C. Patterns of animal migration In: Hansson, L-A, Akesson, S (eds) Animal movement across scales. 1st ed Oxford: Oxford University Press pp 11–30. 2014.

21. Thornton D, Zeller K, Rondinini C, Boitani L, Crooks KR, Burdett C, Rabinowitz A, Quigley H. Assessing the umbrella value of a range-wide conservation network for jaguars (*Panthera onca*). Ecological Applications 2015; 26: 1112–1124.

22. Paviolo A, De Angelo C, Ferraz KMPMB, Morato RG, Pardo JM, et al. A biodiversity hotspot losing its top predator: The challenge of jaguar conservation in the Atlantic Forest of South America Scientific Reports 2016; 6: 37147. DOI: 101038/srep37147.

23. Bogoni JA, Peres CA, Ferraz KMPMB. Extent, intensity and drivers of mammal defaunation: a continental-scale analysis across the Neotropics. Scientific Reports 2020; 10: 14750.

24. Tobler MW, Carillo-Perscastegui SE, Hartley AZ, Powell GVN. High jaguar densities and large population sizes in the core habitat of the southwestern Amazon. Biological Conservation 2013; 159: 375–381

25. Jędrzejewski W, Robinson HS, Abarca M, Zeller KA, Velasquez G, Paemelaere EAD, et al. Estimating large carnivore populations at global scale based on spatial predictions of density and distribution: Application to the jaguar (*Panthera onca*). PLoS ONE 2018; 13(3): e0194719.

26. Eva HD, Huber O, Achard F, Balslev H, Beck S, et al. A proposal for defining the geographical boundaries of Amazonia; synthesis of the results from an expert consultation workshop organized by the European Commission in collaboration with the Amazon Cooperation Treaty Organization-JRC Ispra, 7–8 June 2005 (No 21808- EN). 2005.

27. Nepstad DC, Stickler CM, Soares-Filho B, Merry F, Nin E. Interactions among Amazon land use, forests and climate: prospects for a near-term forest tipping point. Philosophical Transactions of the Royal Society B 2008; 363: 1737–1746.

28. Marques AAB, Schneider M, Peres CA. Human population and socioeconomic modulators of conservation performance in 788 Amazonian and Atlantic Forest reserves PeerJ 4, 2016; pe2206,

29. Simberloff D. Flagships, umbrellas, and keystones: is single-species management passe’ in the landscape era. Biological Conservation 1998; 83: 247–57.

30. Silvério DV, Brando PM, Balch JK, Putz FE, Nepstad DC, Oliveira-Santos C, Bustamante MMC. Testing the Amazon savannization hypothesis: fire effects on invasion of a neotropical forest by native Cerrado and exotic pasture grasses Philosophical Transactions of the Royal Society B 2013; 368: 20120427.

31. Sales LP, Galetti M, Pires MM. Climate and land‐use change will lead to a faunal “savannization” on tropical rainforests Global Change Biology, 2020. DOI:101111/gcb15374.

32. Brazil’s National Institute for Space Research (INPE) Monitoramento do Desmatamento da Floresta Amazônica Brasileira por Satélite Available at: http://www.obtinpebr/OBT/assuntos/programas/amazonia/prodes Access: 28 November 2020. 2020a

33. Brazil’s National Institute for Space Research (INPE). Banco de dados de Queimadas INPE – Programa Queimadas Available at: http://queimadasdgiinpebr/queimadas/bdqueimadas Access: 08 December 2020. 2020b.

34. Silva-Jr CHL, Pessôa ACM, Carvalho NS, Reis JBC, Anderson LO, Aragão LEO. The Brazilian Amazon deforestation rate in 2020 is the greatest of the decade. Nature Ecology and Evolution, 2020. DOI: https://doiorg/101038/s41559-020-01368-x.

35. Walker R, Moore NJ, Arima E, Perz S, et al. Protecting the Amazon with protected areas. Proceedings of the National Academy of Sciences 2009; 106(26): 10582–10586.

36. Gray CL, Hill SLL, Newbold T, Hudson LN, Börger L, Contu S, Hoskins AJ, Ferrier S, Purvis A, Scharlemann JPH. Local biodiversity is higher inside than outside terrestrial protected areas worldwide. Nature Communications 2016; 7: 12306.

37. Begotti RA, Peres CA. Rapidly escalating threats to the biodiversity and ethnocultural capital of Brazilian Indigenous Lands. Land Use Policy 2020; 96: 104694.

38. Walker WS, Gorelik SR, Baccini A, Aragon-Osejo JL, et al. The role of forest conversion, degradation, and disturbance in the carbon dynamics of Amazon indigenous territories and protected areas. Proceedings of the National Academy of Sciences 2020; 117(6): 3015–3025.

39. Moilanen A, Arponen A, Stokland JN, Cabeza M. Assessing replacement cost of conservation areas: How does habitat loss influence priorities? Biological Conservation 2009; 142: 575–585.

40. Almeida-Rocha JA, Peres CA. Nominally protected buffer zones around tropical protected areas are as highly degraded as the wider unprotected countryside Biological Conservation 2021; 256: 109068.

41. Nobre CA, Sampaio G, Borma LS, Catilla-Rubio JC, Silva JS, Cardoso M. Land-use and climate change risks in the Amazon and the need of a novel sustainable development paradigm. Proceedings of the National Academy of Sciences 2016; 113(39): 10759–10768.

42. Kauano EE, Silva JMC, Michalski F. Illegal use of natural resources in federal protected areas of the Brazilian Amazon. PeerJ 2017; 5: e3902, DOI: 107717/peerj3902.

43. Olson DM, Dinerstein E, Wikramanayake ED, Burgess ND, et al. Terrestrial ecoregions of the world: a new map of life on Earth. Bioscience 2011; 51(11): 933–938.

44. Instituto Brasileiro de Geografia e Estatística (IBGE). Censo demográfico Rio de Janeiro Available at: http://www.ibgegovbr Access: 24 April 2021. 2020.

45. Instituto Brasileiro de Geografia e Estatística (IBGE). Censo demogréfico Rio de Janeiro Available at: http://www.ibgegovbr Access: 8 November 2020. 2010.

46. Instituto Brasileiro de Geografia e Estatística (IBGE). BC250 - Base Cartogréfica Contínua do Brasil - 1:250 000 – 2017 Diretoria de Geociências - DGC / Coordenação de Cartografia – CCAR Available at: http://www.metadadosgeoibgegovbr/geonetwork_ibge/srv/por/metadatashow?uuid=5a47e9ea-e2cd-423b-8646-53f67ff4ed2d Access: 10 December 2020. 2017.

47. MapBiomas. Projeto MapBiomas Coleção 5 da Série Anual de Mapas de Cobertura e Uso de Solo do Brasil Available at: https://mapbiomasorg/colecoes-mapbiomas-1?cama_set_language=pt-BR Access: 15 November 2020. 2019.

48. Woodroffe R, Ginsberg JR. Edge effects and the extinction of populations inside protected areas. Science 1998; 280: 2126–2128.

49. ESRI. ArcGIS Desktop: Release 10 Redlands, CA: Environmental Systems Research Institute. 2019.

50. Sistema Nacional de Unidades de Conservação (SNUC). Lei 9985 de 18 de julho de 2000; Ministério do Meio Ambiente. 2000.

51. Ministério do Meio Ambiente (MMA). Cadastro Nacional de Unidades de Conservação (CNUC) Available at: https://antigommagovbr/areas-protegidas/cadastro-nacional-de-ucs/dados-georreferenciadoshtml Access: 25 December 2020. 2019.

52. Fundação Nacional do Índio (FUNAI). Modalidades de Terra Indígenas Available at: http://www.funaigovbr/indexphp/indios-no-brasil/terras-indigenas Access: 1 December 2020. 2019.

53. Zar JH. Biostatistical Analysis 4st ed, New Jersey: Pretince-Hall. 1999.

54. Grace JB. Structural Equation Modeling and Natural Systems. Cambridge University Press. 2006.

55. Dobson AJ. An Introduction to Generalized Linear Models. Chapman and Hall. 1990.

56. Kamata A, Bauer DJ. A Note on the Relation between Factor Analytic and Item Response Theory Models. Structural Equation Modeling 2008; 15(1): 136–153.

57. R Core Team. R: A language and environment for statistical computing R Foundation for Statistical Computing. 2020.

58. Rosseel Y. lavaan: An R Package for Structural Equation Modeling Journal of Statistical Software 2012; 48(2): 1–36.

59. Terborgh J. The role of felid predators in Neotropical Forests. Vida Silvestre Neotropical 1990; 2: 3–5.

60. Laurance WF, Croes BM, Tchingnoumba L, Lahm S, et al. Impacts of roads and hunting on central African rainforest mammals. Conservation Biology 2006; 20(4): 1251–1261.

61. Brando PM, Soares-Filho B, Rodrigues L, Assunção A, et al. The gathering firestorm in southern Amazonia. Science Advances 2020; 6(2): eaay 1632.

62. Ceddia MG, Bardsley NO, Gomez-y-Paloma S, Sedlacek S. Governance, agricultural intensification, and land sparing in tropical South America. Proceedings of the National Academy of Sciences 2014; 111(2): 7242–7247.

63. Brancalion PHS, Garcia LC, Loyola R, Rodrigues RR, Pillar VD, Lewinsohn TN. Análise crítica da Lei de Proteção da Vegetação Nativa (2012), que substituiu o antigo Código Florestal: atualizações e ações em curso. Natureza & Conservação 2016; 14: 1–16.

64. Wilkie DS, Bennett EL, Peres CA, Cunningham AA. The empty forest revisited. Annals of the New York Academy of Sciences 2011; 1223: 120–128.

65. Ferrante L, Fearnside PM. Brazil’s new president and ‘ruralists’ threaten Amazonia’s environment, traditional peoples and the global climate. Environmental Conservation 2019; 46(4): 261–263.

66. Aragão LEOC, Shimabukuro YE. The Incidence of Fire in Amazonian Forests with Implications for REDD. Science 2010; 328: 1275–1278.

67. Barlow J, Peres CA. Fire-mediated dieback and compositional cascade in an Amazonian forest. Philosophical Transactions of the Royal Society B 2008; 363: 1787.

68. Michalski F, Boulhosa RLP, Faria A, Peres CA. Human–wildlife conflicts in a fragmented Amazonian forest landscape: determinants of large felid depredation on livestock. Animal Conservation 2006; DOI: 101111/j1469-1795200600025x.

69. Jorge MLSP, Galetti M, Ribeiro MC, Ferraz KMPMB. Mammal defaunation as surrogate of trophic cascades in a biodiversity hotspot. Biological Conservation 2013; 163: 49–57.

70. Menezes JFS, Tortato FR, Roque FO, Oliveira-Santos LG, Morato RG. Deforestation, fires, and lack of governance are displacing thousands of jaguars in Brazilian Amazon (submitted to Conservation Science and Practice)

71. Morato RG, Connette GM, Stabach JA, De Paula RC, Ferraz KMPM, et al. Resource selection in an apex predator and variation in response to local landscape characteristics. Biological Conservation 2018; 228,: 233–240.

72. Romero-Muñoz A, Torres R, Noss AJ, Giordano AJ, et al. Habitat loss and overhunting synergistically drive the extirpation of jaguars from the Gran Chaco Diversity and Distributions 2018; 1–15.

73. Romero-Muñoz A, Morato RG, Tortato F, Kuemmerle T. Beyond fangs: beef and soybean trade drive jaguar extinction. Frontiers in Ecology and the Environment 2020; 18: 67–68.

74. Vilela T, Harb AM, Bruner A, Arruda VLS, et al. A better Amazon road network for people and the environment. Proceedings of the National Academy of Sciences 2020; 117(13): 7095–7102.

75. Benítez-López A, Alkemade R, Verweij PA. The impacts of roads and other infrastructure on mammal and bird populations: A meta-analysis. Biological Conservation 2010; 143: 1307–1316.

76. Carter N, Killion A, Easter T, Brandt J, Ford A. Road development in Asia: Assessing the range-wide risks to tigers. Science Advances 2020; 6(18): eaaz9619.

77. Joshi AR, Dinerstein E, Wikramanayak E, Anderson ML, et al. Tracking changes and preventing loss in critical tiger habitat. Science Advances 2016; 2: e1501675.

78. Peres CA, Terborgh J. Amazonian Nature Reserves: An Analysis of the Defensibility Status of Existing Conservation Units and Design Criteria for the Future. Conservation Biology 1995; 9(1): 34–46.

79. Stocks A. Too much for too few: problems of indigenous land rights in Latin America Annual. Review of Anthropology 2005; 34: 85–104.

80. Mooers AØ, Faith DP, Maddison WP. Converting Endangered Species Categories to Probabilities of Extinction for Phylogenetic Conservation Prioritization. PLoS ONE 2008; 3(11): e3700.

81. Miranda EBP, Peres CA, Carvalho-Rocha V, Miguel BV, et al. Tropical deforestation induces thresholds of reproductive viability and habitat suitability in Earth’s largest eagles Scientific Reports, In press.

82. Wilson KA, Carwardine J, Possingham HP. Setting Conservation Priorities. Annals of the New York Academy of Sciences 2009; 1162: 237–264.

83. Venter O, Sanderson EW, Magrach A, et al. Sixteen years of change in the global terrestrial human footprint and implications for biodiversity conservation. Nature Communications 2016; 7: 12558.

84. da Silva JMC, Dias TCAC, da Cunha AC, Cunha HFA. Funding deficits of protected areas in Brazil. Land Use Policy 2021; 100: 104926.

